# Inhibition of NF-κB signaling pathway in astrocytes facilitates amyloid-β clearance by kallikrein-related peptidase 7

**DOI:** 10.1101/2025.03.02.641088

**Authors:** Yuki Sudo, Chia-Jen Sung, Kazunori Kikuchi, Yung-Wen Chiu, Sho Takatori, Yukiko Hori, Taisuke Tomita

## Abstract

Alzheimer disease (AD) is characterized by the deposition of amyloid-β peptide (Aβ). Decreased Aβ clearance is observed in sporadic AD patients, suggesting that enhancing Aβ clearance is a potential therapeutic approach for AD. We identified kallikrein-related peptidase 7 (KLK7) as an astrocyte-derived Aβ-degrading protease, and its mRNA expression is reduced in AD brains. Memantine, an N-methyl-D-aspartate (NMDA) receptor antagonist, upregulates *KLK7* expression in astrocytes; however, the regulatory mechanism remains unclear. Here, we show that the NMDA receptor signaling negatively regulates *KLK7* mRNA expression via nuclear factor-κB (NF-κB). Inhibition of NF-κB signaling pathway in astrocytes increases *KLK7* expression and promotes Aβ degradation. Moreover, the mRNA expression level of the NF-κB family is elevated in AD brains and shows a negative correlation with *KLK7* mRNA expression. Finally, the injection of an NF-κB inhibitor significantly upregulates *Klk7* expression and reduces Aβ levels *in vivo*. These findings suggest that the NMDA receptor-NF-κB signaling axis in astrocytes negatively regulates *KLK7* expression and modulates KLK7-mediated Aβ clearance.

## Introduction

Alzheimer disease (AD) is a progressive neurodegenerative disorder and the most common form of dementia^1^. The pathological hallmark of AD is the deposition of amyloid-β peptide (Aβ) as senile plaques in the brain. Aβ is produced through the proteolytic cleavage of amyloid precursor protein (APP) in neurons^2^. Genetic mutations found in familial AD patients increase Aβ production and aggregation^3–5^. However, sporadic AD patients, accounting for more than 90% of AD cases, exhibit a decreased rate of Aβ clearance rather than an increased rate of Aβ production^6^. Thus, understanding the molecular mechanisms of Aβ clearance is considered necessary for the development of AD treatments.

Degradation of Aβ by proteases is one of the mechanisms for Aβ clearance^7^. Neprilysin (NEP), a major Aβ-degrading protease, is primarily expressed in neurons^8^. However, since neurons degenerate in AD brains, other cell types such as glial cells might serve as more suitable targets for therapeutic approaches^9^. Recently, the role of astrocytes in AD has been highlighted. Astrocytes are the most abundant glial cells, which perform diverse functions in maintaining the brain environment^10–12^. Astrocytes become reactive in response to brain injury and diseases such as AD^13^. Various central nervous system (CNS) stimuli, like cytokines, induce several types of reactive astrocytes with different functions and transcriptomes^14^. For example, a neurotoxic type of reactive astrocytes termed A1 astrocytes is destructive to synapses^14^, characterized by the activation of nuclear factor-κB (NF-κB) signaling pathway^15^. In addition, a type of reactive astrocytes specifically appearing in early disease stages of an AD mouse model has been identified, with enriched expression of genes involved in endocytosis, complement cascade, and aging^16^. These findings suggest the heterogeneity of reactive astrocytes. However, the pathological role of astrocytes in AD, especially the contributions of the signaling pathways activated in reactive astrocytes to Aβ clearance, remains to be elucidated.

We recently reported that the conditioned medium from astrocytes exhibits robust Aβ-degrading activity and that kallikrein-related peptidase 7 (KLK7) is responsible for this activity^17^. *KLK7* mRNA expression is significantly decreased in AD brains, and knockout of *Klk7* exacerbates Aβ deposition in AD model mice, highlighting the critical role of KLK7 in Aβ pathology. In addition, our findings, corroborated by an independent study, have shown that KLK7 is capable of degrading not only Aβ monomers but also Aβ fibrils, thereby reducing cell toxicity *in vitro*^17,18^. Intriguingly, memantine, an N-methyl-D-aspartate (NMDA) receptor antagonist used in AD treatment, upregulates *KLK7* expression in astrocytes. However, the underlying mechanism remained unclear.

In this study, we investigated the mechanism of *KLK7* expression regulated by NMDA receptor signaling. Using a promoter reporter system, we found that NMDA receptor signaling downregulates *KLK7* transcription, and that NF-κB is involved in the transcriptional repression. Moreover, in AD patients with decreased *KLK7* mRNA expression, the expressions of multiple NF-κB family members are significantly increased. Finally, pharmacological inhibition of the NF-κB signaling ameliorates Aβ pathology in the brains of AD model mice, suggesting that NF-κB functionally regulates the Aβ-degrading capacity of KLK7 *in vivo*.

## Results

### NMDA receptor signaling negatively regulates KLK7 mRNA expression and Aβ degradation

Our group previously demonstrated that anti-dementia drug memantine, an NMDA receptor antagonist, increases *Klk7* expression in mouse primary astrocytes^17^. In this study, we used the neuroglioma cell line H4 for further investigation. H4 cells were treated with memantine for 24 hours, and *KLK7* mRNA expression was assessed by using quantitative reverse transcription PCR (qRT-PCR). The results showed a significant increase in *KLK7* mRNA expression (**Fig. 1a**). In contrast, NMDA or L-glutamate treatment significantly downregulated *KLK7* expression, indicating the negative regulation of *KLK7* expression by NMDA receptor signaling in H4 cells (**Fig. 1b**). Note that, the culture medium we used likely contains small amounts of L-glutamate, which can be generated from L-glutamine. This presence of L-glutamate may result in a low-level suppression of *KLK7* expression. Next, we tested the effects of memantine on the Aβ-degrading activity of H4 cells. Synthetic Aβ40 and memantine were co-treated, and the conditioned medium was collected after 24 hours (**Fig. 1c**). Remaining Aβ40 in the conditioned medium was then evaluated by immunoblotting analysis. The results showed that memantine enhanced Aβ-degrading activity of H4 cells without affecting cell viability (**Fig. 1d, e**). These findings suggest that inhibition of the NMDA receptor signaling upregulates *KLK7* mRNA expression and promotes Aβ degradation.

**Fig. 1.**
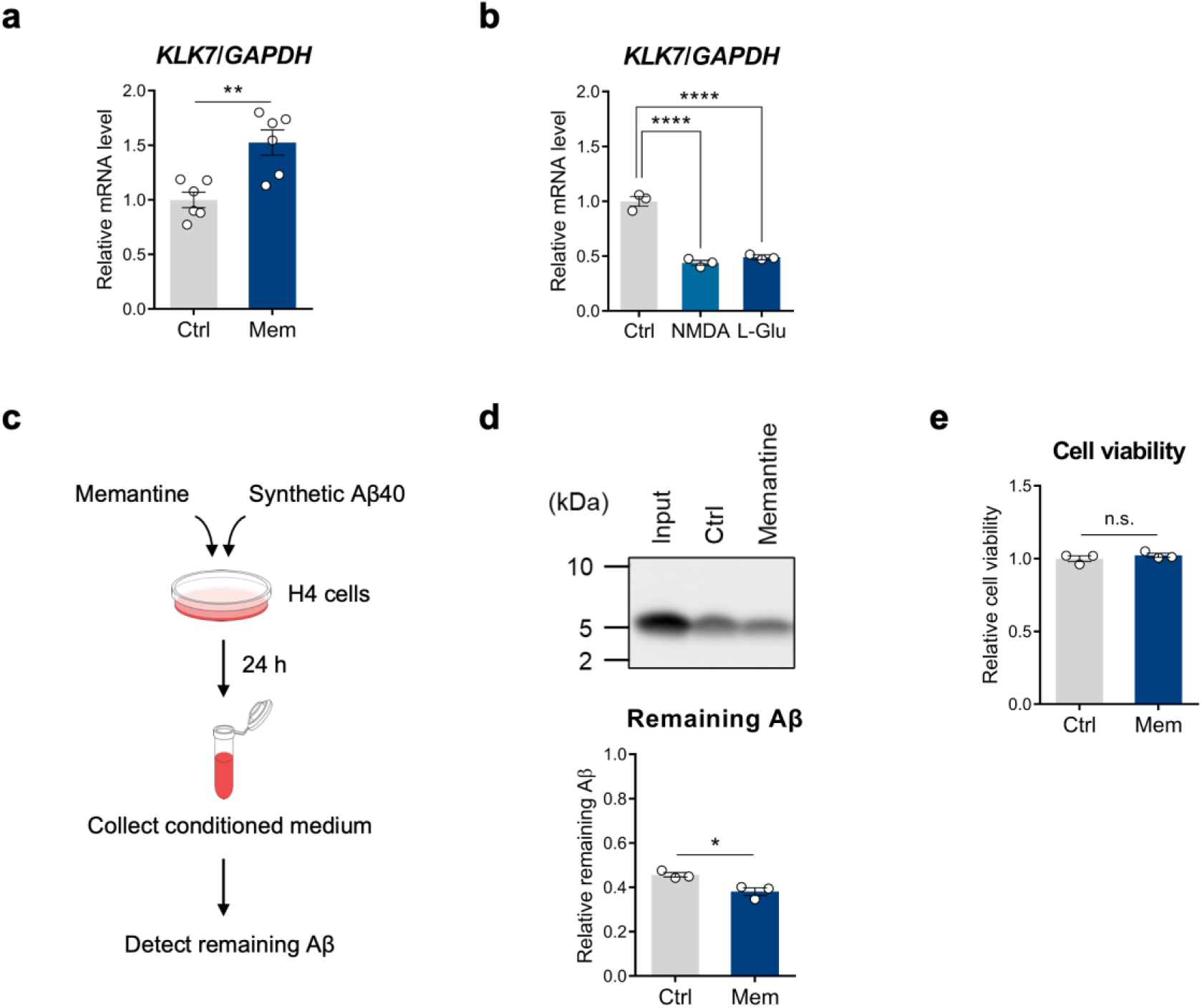
NMDA receptor signaling negatively regulates *KLK7* mRNA expression and Aβ degradation. **a, b** The mRNA expression level of *KLK7* in H4 cells treated with 30 µM memantine (Mem) (n=6, Student’s *t*-test, p=0.003) **(a)**, 100 µM NMDA, and 100 µM L-glutamate (L-Glu) (n=3, Dunnett’s test, p_Ctrl vs NMDA_<0.0001, p_Ctrl vs L-Glu_<0.0001) **(b)**, respectively. **c** Schematic representation of Aβ degradation assay in H4 cells. Synthetic Aβ40 was added to the conditioned medium of H4 cells treated with memantine. After 24 hours, remaining Aβ40 in conditioned medium was detected by immunoblotting. **d** The Aβ-degrading activity of H4 cells treated with 30 µM memantine. A representative immunoblot and quantified results are shown (Input=1.0, n=3, Student’s *t*-test, p=0.0205). **e** Cell viability of H4 cells treated with 30 µM memantine (n=3, Student’s *t*-test, p=0.3927). All data are shown as mean±SEM. *p<0.05, **p<0.01, ****p<0.0001; n.s., non-significant.

### NF-κB is involved in the NMDA receptor signaling-mediated transcriptional repression of KLK7

Next, to investigate the mechanism by which NMDA receptor signaling regulates *KLK7* expression, we analyzed the transcriptional activity of *KLK7* using a luciferase reporter assay. We generated luciferase constructs containing the 238-bp promoter region upstream of the human *KLK7* gene^19^. Luciferase constructs were introduced into H4 cells to establish stable cell lines, termed H4 h238 WT, with H4 Empty, which had no promoter sequence, serving as a control (**Fig. 2a**). To verify the functionality of the luciferase reporters, we treated H4 stable cells with Trichostatin A (TSA), a histone deacetylase (HDAC) inhibitor known to upregulate transcription. Luciferase activity of H4 h238 WT was enhanced by TSA treatment, while that of H4 Empty remained unchanged as expected (**Supplementary Fig. 1a**). We then treated H4 h238 WT with NMDA or L-glutamate and collected the cells 24 hours later for luciferase reporter assay. Luciferase activity of H4 h238 WT was significantly reduced upon treatment with NMDA and L-glutamate (**Fig. 2b**). Conversely, memantine treatment enhanced the luciferase activity of H4 h238 WT (**Fig. 2c**). These results suggest that NMDA receptor signaling downregulates the transcriptional activity of *KLK7*.

**Fig. 2.**
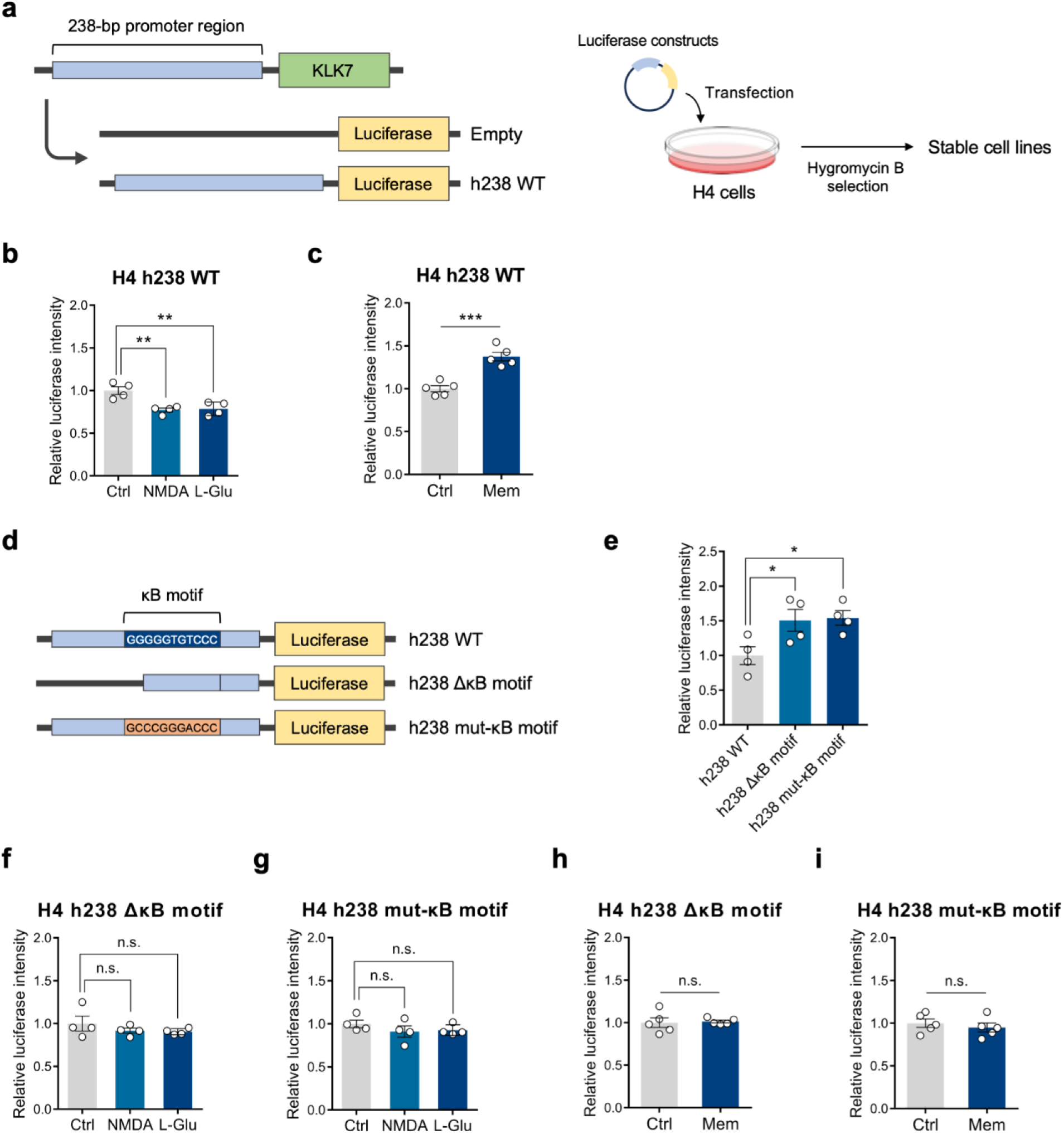
NF-κB is involved in the NMDA receptor signaling-mediated transcriptional repression of *KLK7*. **a** Schematic representation of the establishment of luciferase constructs (Empty and h238 WT) and H4 stable cell lines. **b, c** Luciferase activity of H4 h238 WT treated with 100 µM NMDA, 100 µM L-glutamate (n=4, Dunnett’s test, p_Ctrl vs NMDA_=0.0038, p_Ctrl vs L-Glu_=0.0057) **(b)**, and 30 µM memantine (n=5, Student’s *t*-test, p=0.0003) **(c)**, respectively. **d** Schematic representation of luciferase constructs containing a deletion mutant and a motif mutant of the κB motif in the 238-bp *KLK7* promoter region (h238 ΔκB motif and h238 mut-κB motif). **e** Luciferase activity of each luciferase construct transiently expressed in H4 cells (n=4, Dunnett’s test, p_h238 WT vs h238 ΔκB motif_=0.0432, p_h238 WT vs h238 mut-κB motif_=0.0318). **f, g** Luciferase activity of H4 h238 ΔκB motif (n=4, Dunnett’s test, p_Ctrl vs NMDA_=0.4847, p_Ctrl vs L-Glu_=0.3969) **(f)** and H4 h238 mut-κB motif (n=4, Dunnett’s test, p_Ctrl vs NMDA_=0.3716, p_Ctrl vs L-Glu_=0.5012) **(g)** treated with 100 µM NMDA or 100 µM L-glutamate. **h, i** Luciferase activity of H4 h238 ΔκB motif (n=5, Student’s *t*-test, p=0.8164) **(h)** and H4 h238 mut-κB motif (n=5, Student’s *t*-test, p=0.5011) **(i)** treated with 30 µM memantine. All data are shown as mean±SEM. *p<0.05, **p<0.01, ***p<0.001; n.s., non-significant.

Several transcription factors were predicted to bind to the 238-bp promoter region of the *KLK7* gene^19^. Among these, we focused on the involvement of NF-κB. NF-κB is a family of transcription factors that play crucial roles in immune responses and inflammation^20^. In mammals, the NF-κB family consists of five proteins: p65, RelB, c-Rel, p50, and p52. These proteins form dimers with each other to regulate gene expression^20^. NF-κB is activated in response to various stimuli, including cytokines, growth factors, and glutamate in the CNS^21^. Given its reported significance in reactive astrocytes^14,15^, we examined the role of NF-κB in regulating *KLK7* expression and its potential contribution to the Aβ clearance ability of astrocytes. NF-κB proteins bind to a consensus sequence known as the κB motif^22^ (**Supplementary Fig. 2a**). The potential κB motif was confirmed to be present in the 238-bp *KLK7* promoter region (**Supplementary Fig. 2b**). We generated two luciferase constructs: one containing a deletion mutant of the κB motif (h238 ΔκB motif) and the other containing a motif mutant of the κB motif (h238 mut-κB motif) in the 238-bp *KLK7* promoter region (**Fig. 2d**). We transiently expressed the h238 ΔκB motif and h238 mut-κB motif in H4 cells and performed a luciferase reporter assay. Luciferase activity of h238 ΔκB motif and h238 mut-κB motif was significantly increased compared to that of h238 WT, suggesting the inhibitory role of NF-κB in *KLK7* transcription (**Fig. 2e**). Next, to determine whether NF-κB functions downstream of the NMDA receptor signaling, we established stable cell lines (H4 h238 ΔκB motif and H4 h238 mut-κB motif) and treated them with NMDA or L-glutamate. The results demonstrated that both H4 h238 ΔκB motif and H4 h238 mut-κB motif mitigated the suppression of *KLK7* transcriptional activity caused by NMDA and L-glutamate, suggesting the involvement of NF-κB in the regulatory pathway modulated by NMDA receptor signaling (**Fig. 2f, g**). Moreover, both H4 h238 ΔκB motif and H4 h238 mut-κB motif attenuated the enhancement of luciferase activity induced by memantine (**Fig. 2h, i**). These results collectively indicate that NF-κB is involved in the NMDA receptor signaling-mediated negative regulation of *KLK7* transcription.

### Inhibition of the NF-κB signaling pathway increases KLK7 mRNA expression and promotes Aβ degradation

NF-κB signaling is activated through either the canonical or the noncanonical pathway. The canonical pathway drives the expression of genes essential for inflammatory response, cell proliferation, and survival, while the noncanonical pathway primarily regulates the development of immune cells and lymphoid organs^23,24^. In the canonical pathway, NF-κB proteins are initially bound and inhibited by the IκB proteins. Upon stimulation by factors such as proinflammatory cytokines and growth factors, the IκB kinase (IKK) complex is activated and phosphorylates the IκB proteins. Phosphorylated IκB proteins undergo subsequent ubiquitination and degradation via the proteasome, resulting in the release of NF-κB proteins. This process exposes the nuclear translocation signal sequence of NF-κB proteins, allowing them to translocate to the nucleus and regulate gene expression^21^ (**Fig. 3a**). In contrast, the noncanonical pathway involves IKKα-mediated phosphorylation of p100, the precursor of p52. Phosphorylated p100 undergoes proteasomal processing to generate p52, enabling the formation of p52/RelB complex that translocates to the nucleus^21,23^.

**Fig. 3.**
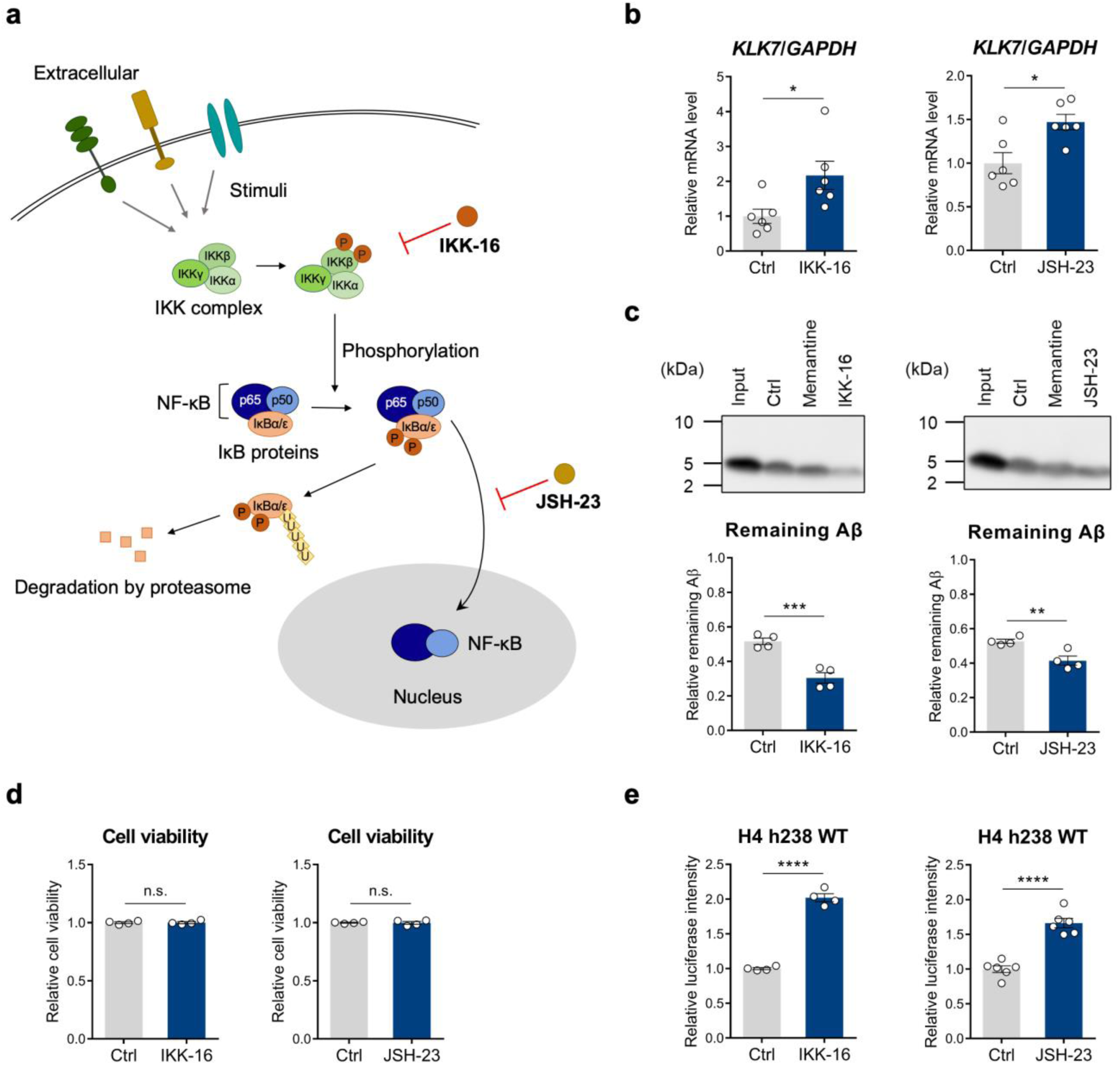
Inhibition of the NF-κB signaling pathway increases *KLK7* mRNA expression and promotes Aβ degradation. **a** Schematic representation of the canonical NF-κB activation pathway with the targets of the inhibitors IKK-16 and JSH-23 indicated. **b** The mRNA expression level of *KLK7* in H4 cells treated with 2 µM IKK-16 or 30 µM JSH-23 (n=6, Student’s *t*-test, p_Ctrl vs IKK-16_=0.0270, p_Ctrl vs JSH-23_=0.0103). **c** The Aβ-degrading activity of H4 cells treated with 2 µM IKK-16 or 30 µM JSH-23. Representative immunoblots and quantified results are shown (Input=1.0, n=4, Student’s *t*-test, p_Ctrl vs IKK-16_=0.0010, p_Ctrl vs JSH-23_=0.0082). **d** Cell viability of H4 cells treated with 2 µM IKK-16 or 30 µM JSH-23 (n=4, Student’s *t*-test, p_Ctrl vs IKK-16_=0.9618, p_Ctrl vs JSH-23_=0.9043). **e** Luciferase activity of H4 h238 WT treated with 2 µM IKK-16 or 30 µM JSH-23 (n=4 (IKK-16), n=6 (JSH-23), Student’s *t*-test, p_Ctrl vs IKK-16_<0.0001, p_Ctrl vs JSH-23_<0.0001). All data are shown as mean±SEM. *p<0.05, **p<0.01, ***p<0.001, ****p<0.0001; n.s., non-significant.

To investigate the role of NF-κB in the regulation of *KLK7* expression, we used two inhibitors targeting the NF-κB signaling pathway. The first inhibitor, IKK-16, inhibits the IKK complex, which mediates NF-κB activation^25^. The second inhibitor, JSH-23, inhibits NF-κB nuclear translocation^26^ (**Fig. 3a**). Treatment of H4 cells with IKK-16 or JSH-23 for 24 hours resulted in an increase in *KLK7* mRNA expression (**Fig. 3b**). Analysis of Aβ-degrading activity in H4 cells revealed a significant increase in Aβ degradation following IKK-16 and JSH-23 treatments (**Fig. 3c**). Importantly, we observed no changes in cell viability (**Fig. 3d**). Similar experiments were conducted using mouse primary glial cultures, which were predominantly composed of astrocytes. Treatment with IKK-16 increased *Klk7* mRNA expression and enhanced Aβ-degrading activity in mouse primary astrocytes without altering cell viability (**Supplementary Fig. 3a-c**). Additionally, luciferase reporter assay demonstrated that both IKK-16 and JSH-23 increased the luciferase activity of H4 h238 WT (**Fig. 3e**). Collectively, these results suggest that inhibition of the NF-κB signaling pathway in astrocytes activates *KLK7* transcription and enhances Aβ-degrading activity.

### The mRNA expressions of the NF-κB family and KLK7 are negatively correlated in AD brains

As mentioned above, NF-κB regulates gene expression by forming homo- or heterodimers, which are composed of proteins from the five NF-κB family members: p65, RelB, c-Rel, p50, and p52, encoded by *RELA*, *RELB*, *REL*, *NFKB1*, and *NFKB2*, respectively^20^. Given our previous findings of decreased *KLK7* mRNA levels in AD brains^17^, we analyzed the mRNA expressions of NF-κB proteins in AD brains and explored their correlation with *KLK7* mRNA expression. We analyzed two public RNA sequencing (RNA-seq) datasets deposited in the Accelerating Medicines Partnership - Alzheimer’s Disease (AMP-AD) Knowledge Portal, namely, the Mayo RNA-seq^27^ and Mount Sinai Brain Bank (MSBB) AD cohorts^28^. In the Mayo RNA-seq dataset, *KLK7* mRNA level was decreased in the temporal cortex of AD patients, as expected. In contrast, the mRNA expression levels of *RELA, NFKB1*, and *NFKB2* were significantly increased (**Fig. 4a**). Similarly, in the MSBB dataset, *KLK7* mRNA expression was reduced, while *RELA*, *NFKB1*, and *NFKB2* mRNA expressions were significantly elevated in the parahippocampal gyrus (Brodmann area 36) of AD brains (**Fig. 4b-g**). Furthermore, the mRNA expression levels of *RELA, REL, NFKB1*, and *NFKB2* were negatively correlated with *KLK7* mRNA expression (**Fig. 4h-l**). These findings collectively suggest that the NF-κB signaling pathway is activated in AD brains which may contribute to the decreased expression of *KLK7*.

**Fig. 4.**
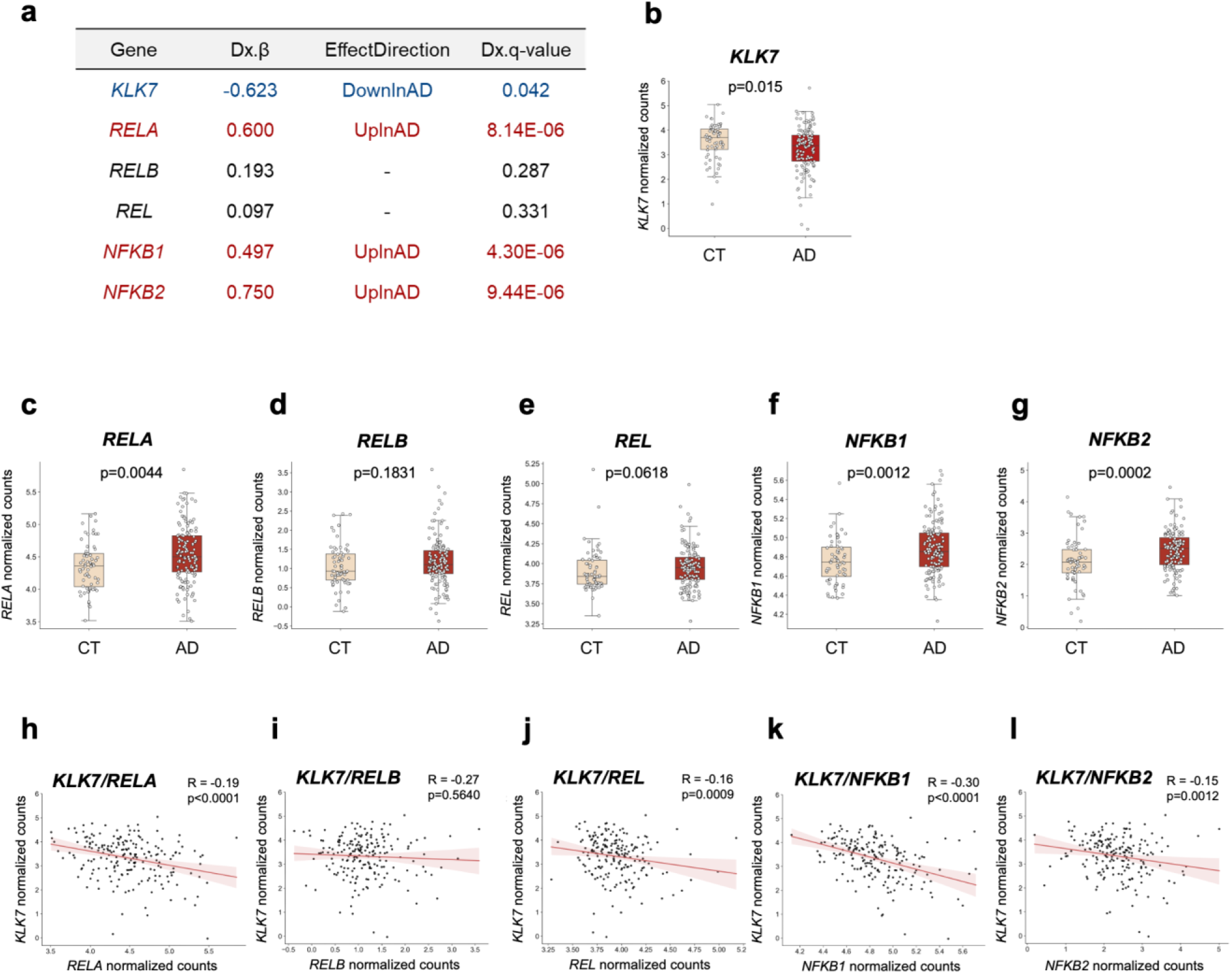
The mRNA expressions of the NF-κB family and *KLK7* are negatively correlated in AD brains. **a** Comparison of *KLK7*, *RELA*, *RELB*, *REL*, *NFKB1*, and *NFKB2* mRNA levels between control (n=80) and AD (n=84) subjects from Mayo RNA-seq AD cohort are shown. Dx.β represents the regression coefficient. Effect direction indicates the changes in target mRNA levels between control and AD subjects. Dx.q-value represents the adjusted p-value, with significant differences defined as Dx.q-value<0.05. **b–g** Changes in the mRNA expression levels of *KLK7* **(b)**, *RELA* **(c)**, *RELB* **(d)**, *REL* **(e)**, *NFKB1* **(f)**, and *NFKB2* **(g)** in the brains of healthy control subjects (CT) (n=64) and AD patients (n=137) from MSBB AD cohort are shown. p-values were assessed using the Mann-Whitney *U* test. **h–l** Correlations of mRNA expressions between *KLK7* and *RELA* **(h)**, *RELB* **(i)**, *REL* **(j)**, *NFKB1* **(k)**, and *NFKB2* **(l)** in the brains of 201 subjects from MSBB AD cohort are shown. The regression line is indicated as the red line. Light red background indicates the confidence interval. The correlation coefficients and p-values were assessed using the Kendall rank correlation coefficient.

### Inhibition of the NF-κB signaling pathway increases Klk7 mRNA expression and ameliorates Aβ pathology in vivo

Finally, we investigated the role of NF-κB in regulating *Klk7* expression and Aβ pathology *in vivo*. Given that IKK-16 demonstrated a stronger effect on both *KLK7* mRNA expression and Aβ-degrading activity than JSH-23 *in vitro* (**Fig. 3b, c**), we selected IKK-16 for *in vivo* experiments. IKK-16 was injected into the left hippocampus of 7-month-old wild-type male mice, while the right hippocampus received a vehicle injection as a control (**Fig. 5a**). Twenty-four hours after injection, hippocampal samples were collected for qRT-PCR analysis. The results showed that *Klk7* mRNA expression was significantly increased in the hippocampal samples following IKK-16 injection (**Fig. 5b**).

**Fig. 5.**
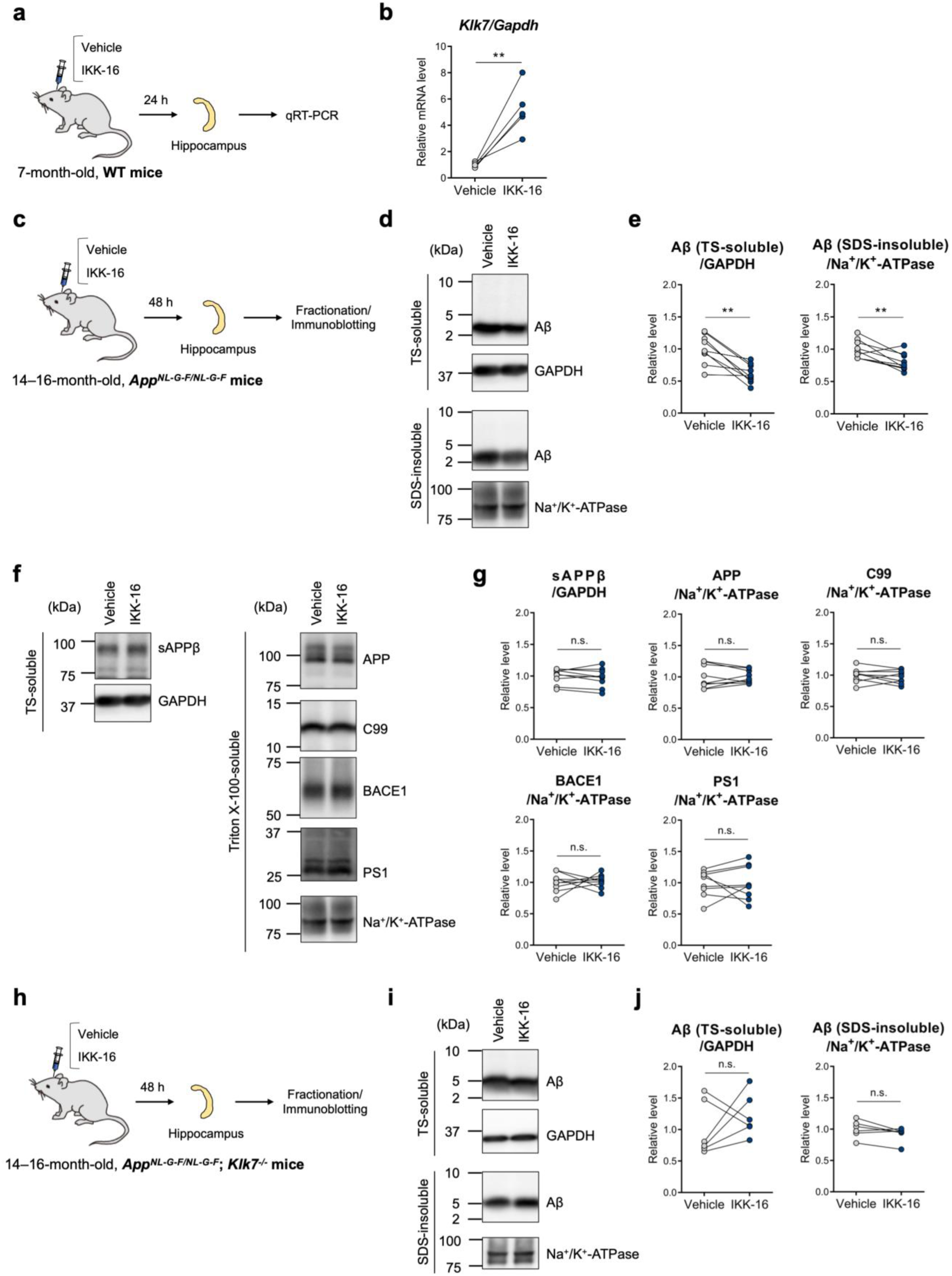
Inhibition of the NF-κB signaling pathway increases *Klk7* mRNA expression and ameliorates Aβ pathology *in vivo*. **a** Schematic representation of IKK-16 injection into the hippocampus of wild-type (WT) mice. **b** The mRNA expression level of *Klk7* in the hippocampus of WT mice injected with 100 µM IKK-16 (n=5, paired *t*-test, p=0.0073). **c** Schematic representation of IKK-16 injection into the hippocampus of *App^NL-G-F/NL-G-F^* mice. **d** Representative immunoblots of TS-soluble Aβ, GAPDH (in the TS-soluble fraction), SDS-insoluble Aβ, and Na^+^/K^+^-ATPase (in the SDS-insoluble fraction) in the hippocampus of *App^NL-G-F/NL-G-F^*mice injected with 100 µM IKK-16. **e** Quantified results of **d** (n=9, paired *t*-test, p_TS-soluble Aβ_=0.0014, p_SDS-insoluble Aβ_=0.0075). **f** Representative immunoblots of sAPPβ, GAPDH (in the TS-soluble fraction), APP, C99, BACE1, PS1, and Na^+^/K^+^-ATPase (in the Triton X-100-soluble fraction) in the hippocampus of *App^NL-G-F/NL-G-F^*mice injected with 100 µM IKK-16. **g** Quantified results of **f** (n=9, paired *t*-test, p_sAPPβ_=0.3201, p_APP_=0.9492, p_C99_=0.4748, p_BACE1_=0.8260, p_PS1_=0.9882). **h** Schematic representation of IKK-16 injection into the hippocampus of *App^NL-G-F/NL-G-F^*; *Klk7^−/−^* mice. **i** Representative immunoblots of TS-soluble Aβ, GAPDH (in the TS-soluble fraction), SDS-insoluble Aβ, and Na^+^/K^+^-ATPase (in the SDS-insoluble fraction) in the hippocampus of *App^NL-G-F/NL-G-F^*; *Klk7^−/−^* mice injected with 100 µM IKK-16. **j** Quantified results of **i** (n=6, paired *t*-test, p_TS-soluble Aβ_=0.3927, p_SDS-insoluble Aβ_=0.1001). **p<0.01; n.s., non-significant.

Next, we investigated the effect of IKK-16 injection on Aβ degradation in *App^NL-G-F/NL-G-F^*mice. *App^NL-G-F/NL-G-F^* mice are human *APP* knock-in models carrying three familial AD mutations in the humanized *APP* sequence: the Swedish (KM670/671NL) mutation, the Beyreuther/Iberian (I716F) mutation, and the Arctic (E22G) mutation^29^. Aβ deposition in the brains of *App^NL-G-F/NL-G-F^* mice begins at 2 months and nearly reaches saturation by 7 months^29^. IKK-16 was injected into the hippocampus of 14–16-month-old female *App^NL-G-F/NL-G-F^* mice (**Fig. 5c**). Forty-eight hours post-injection, hippocampal samples were collected and fractionated into the Tris Saline (TS)-soluble fraction, the Triton X-100-soluble fraction, and the SDS-insoluble fraction. The TS-soluble fraction mainly contains soluble proteins, while the SDS-insoluble fraction is enriched with insoluble proteins, including aggregated Aβ. The Triton X-100-soluble fraction mainly contains membrane proteins. Immunoblotting analysis revealed that Aβ levels in the TS-soluble and SDS-insoluble fractions were significantly decreased in the IKK-16-injected hippocampal samples (**Fig. 5d, e**). The Triton X-100-soluble fraction contained no detectable Aβ. We also examined the effect of IKK-16 on the Aβ production pathway. The protein levels of sAPPβ, APP, C99, β-site APP cleaving enzyme 1 (BACE1), and presenilin 1 (PS1) remained unchanged following IKK-16 injection, suggesting that inhibition of the NF-κB signaling did not affect the Aβ production machinery (**Fig. 5f, g**). Additionally, IKK-16 injection had no effect on the protein levels of insulin-degrading enzyme (IDE) and NEP, the two major Aβ-degrading proteases (**Supplementary Fig. 4a, b**). Moreover, injection of IKK-16 into the hippocampus of *App^NL-G-F/NL-G-F^*; *Klk7*^−/−^ female mice abolished the induced Aβ degradation observed in the hippocampal samples of IKK-16-injected *App^NL-G-F/NL-G-F^*mice (**Fig. 5h-j**). This effect occurred without altering the Aβ production pathway (**Supplementary Fig. 5a, b**). Altogether, these results indicated that NF-κB negatively regulates *Klk7* expression, and inhibition of the NF-κB signaling enhances Aβ degradation mediated by KLK7 *in vivo*.

## Discussion

In this study, we demonstrated that NMDA receptor signaling negatively regulates *KLK7* expression via NF-κB (**Fig. 6a**). Inhibition of the NMDA receptor-NF-κB signaling axis in astrocytes significantly increases *KLK7* expression and augments Aβ-degrading activity (**Fig. 6b**). Moreover, in AD brains, the mRNA expression levels of multiple NF-κB family members are elevated and inversely correlated with *KLK7* mRNA expression. Finally, we revealed that inhibiting the NF-κB signaling pathway in the brains of AD model mice significantly reduces Aβ levels, underscoring the role of NF-κB in regulating KLK7-mediated proteolytic clearance of Aβ *in vivo*.

**Fig. 6.**
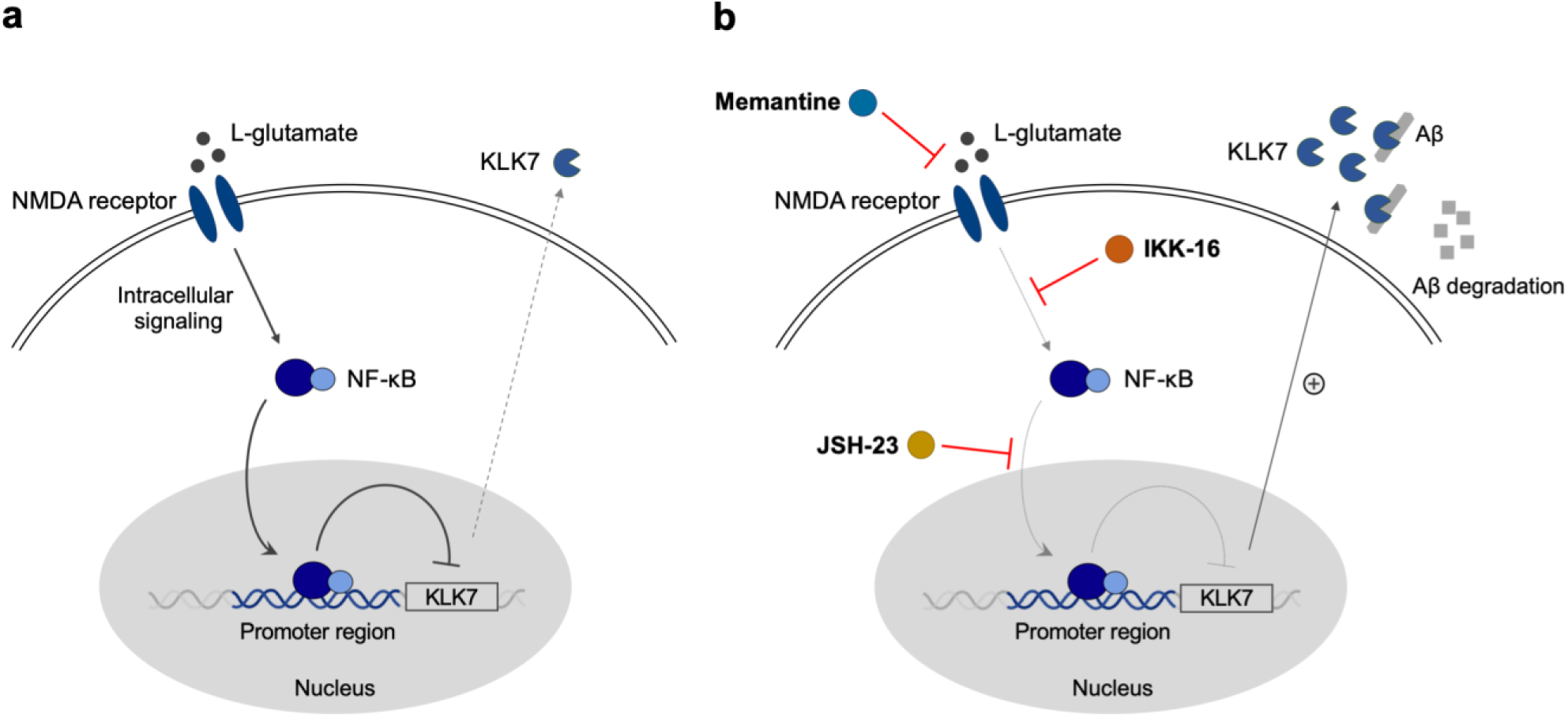
The NMDA receptor-NF-κB signaling axis in astrocytes suppresses *KLK7* expression and modulates KLK7-mediated Aβ clearance. **a** NMDA receptor signaling negatively regulates *KLK7* expression via NF-κB. **b** Inhibition of the NMDA receptor-NF-κB signaling axis by memantine, IKK-16, and JSH-23 upregulates *KLK7* expression and enhances Aβ degradation.

The NMDA receptor is an ionotropic glutamate receptor mediating excitatory synaptic transmission in the CNS^30^. Although primarily expressed in neurons^31^, the NMDA receptor is also found in astrocytes, where it plays a role in neuronal-glial signal transmission^32–34^. In this study, we demonstrated that memantine, an NMDA receptor antagonist, increased *KLK7* expression and enhanced Aβ-degrading activity in a neuroglioma cell line, while treatment with NMDA or L-glutamate led to decreased *KLK7* expression (**Fig. 1**). NMDA receptors typically form tetrameric complexes composed of two GluN1 subunits and two GluN2 subunits, with the GluN2 subunits having four types known as GluN2A, 2B, 2C and 2D^35^. Among these, GluN2C has been identified as a subtype specifically expressed in astrocytes^36,37^, and memantine exhibits higher selectivity for GluN2C/D-containing receptors compared to GluN2A/B-containing receptors^38^. Our previous study demonstrated that MK-801, a non-selective NMDA receptor antagonist, did not induce an increase in *Klk7* expression in mouse primary astrocytes^17^. Therefore, the regulation of *KLK7* expression by NMDA receptor signaling might be linked to the unique sensitivity of astrocytic GluN2C.

Results from the luciferase reporter assay revealed that NMDA receptor signaling suppressed *KLK7* transcription, and this suppression was canceled by the mutation in the κB motif located in the *KLK7* promoter region (**Fig. 2**). These results suggest that NF-κB is involved in the negative regulation of *KLK7* transcription mediated by NMDA receptor signaling. Several studies have indicated that NMDA receptor signaling regulates NF-κB in the CNS. For example, memantine is known to downregulate the HMGB1/TLR4/NF-κB inflammatory axis to ameliorate diabetic neuropathic pain in mice^39^. In addition, it has been reported that memantine attenuates blood-brain barrier damage by suppressing NF-κB-induced cytokine production during cerebral ischemia-reperfusion^40^. Furthermore, the NMDA receptor is known to particularly facilitate the Ca^2+^ influx, and studies have shown that Ca^2+^ signaling activates NF-κB via the PI3K-Akt signaling pathway in granule cells of the cerebellum and in the cortical neurons^41,42^. However, further analyses are required to fully elucidate the detailed pathway between NMDA receptor signaling and NF-κB to clarify the upstream mechanism of *KLK7* expression.

We observed that inhibiting the NF-κB signaling pathway in astrocytes increased *KLK7* expression and enhanced Aβ-degrading activity (**Fig. 3 and Supplementary Fig. 3**). These results indicate that NF-κB negatively regulates *KLK7* expression. Transcriptional repression involving p65, a major NF-κB family member, has been reported for several genes, including ApoE, gastrin, and catechol-O-methyltransferase^43–45^. Although the detailed mechanisms in these cases remain unclear, it is known that p65 interacts with histone-modifying enzymes, and the p65-mediated recruitment of HDAC1 to the gene promoter region may result in transcriptional repression^46^. Moreover, NF-κB family members form homo- or heterodimers, and the functional roles of NF-κB dimers vary among different cell types and target genes^47^. For instance, p65-containing dimers act on the interleukin 8 promoter, while Rel-containing dimers selectively regulate the transcription of interleukin 12-p35 subunits^48^. Additionally, *CXCL13* expression is regulated by p52/RelB, whereas the expression of urokinase plasminogen activator is regulated by p65/Rel selectively^49^. Thus, it is necessary to clarify which NF-κB dimers primarily regulate *KLK7* expression in astrocytes through analyses such as chromatin immunoprecipitation.

We have previously reported that the expression level of *KLK7* mRNA is significantly decreased in the brains of AD patients^17^. In addition, the protein level of KLK7 in the cerebrospinal fluid of AD patients is also reduced^50^. Expression analysis of human brain samples revealed elevated expression levels of multiple NF-κB family members in AD, which were negatively correlated with *KLK7* expression. (**Fig. 4**). These findings suggest that abnormal activation of the NF-κB signaling pathway in AD brains may be associated with the decrease in *KLK7* expression. In fact, NF-κB is activated in A1 astrocytes, a subtype of reactive astrocytes characterized by neurotoxic features in AD^14,15^. Moreover, inhibiting the astrocytic NF-κB signaling pathway can ameliorate neuroinflammation and mitigate neurodegeneration^51^. Additionally, NMDA receptor signaling in astrocytes may be activated in AD brains due to increased glutamate release and decreased glutamate uptake by astrocytes and fibroblasts^52–54^. Nevertheless, genetic analysis and single-cell RNA-seq analysis are needed to further examine the role of the NMDA receptor-NF-κB signaling axis in the transcriptional regulation of *KLK7* in AD astrocytes.

Finally, we explored the potential of NF-κB as a therapeutic target *in vivo*. First, inhibiting the NF-κB signaling pathway by IKK-16 injection into the hippocampus of wild-type mice significantly increased *Klk7* mRNA expression. Next, IKK-16 injection into the hippocampus of *App^NL-G-F/NL-G-F^* mice, but not *App^NL-G-F/NL-G-F^*; *Klk7*^−/−^ mice, reduced Aβ levels without affecting the Aβ production machinery (**Fig. 5**). However, since NF-κB is involved in various aspects of immune function and inflammation^55^, global inhibition of the NF-κB signaling pathway raises concerns about impaired immune responses^56^. Furthermore, NF-κB plays a pivotal role in synaptic transmission, neuronal plasticity, and neuronal development in the CNS^57^. Thus, further investigations are needed to identify cell type-specific and more detailed regulatory mechanisms of *KLK7* expression mediated by NF-κB for developing therapeutic interventions.

Notably, IKK-16 injection into the hippocampus of *App^NL-G-F/NL-G-F^*mice reduced both soluble and insoluble Aβ levels (**Fig. 5d, e**). This may be due to the Aβ-degrading capacity of KLK7, which degrades not only monomeric Aβ but also insoluble Aβ fibrils. Although various proteases can degrade monomeric Aβ, fibrillar Aβ is not targeted by major Aβ-degrading proteases like NEP or IDE^58,59^. Importantly, Aβ fibrils are the main component of amyloid plaques in AD brains, and recently approved anti-Aβ antibodies (i.e., lecanemab and donanemab) specifically remove the aggregated Aβ species. Thus, targeting KLK7 could be a promising strategy for mitigating Aβ pathology in AD. Our findings provide new insights into potential therapeutic approaches for AD by modulating *KLK7* expression in astrocytes through the NMDA receptor-NF-κB signaling axis.

## Methods

### Cell culture

H4 cells were purchased from ATCC. Cells were cultured in Dulbecco’s modified Eagle’s medium (DMEM) with high glucose (Wako) added with 10% heat-inactivated fetal bovine serum (Hyclone), 50 units/mL penicillin (Wako), and 50 unit/mL streptomycin (Wako) at 37°C in humidified air containing 5% CO_2_^17^. Primary glial cells, mainly comprised of astrocytes, were prepared as previously described with several modifications^17,60^. Briefly, the cerebrum was isolated from each mouse at postnatal day 2 (P2) in ice-cold HBSS (−) (Wako) and then suspended in HBSS (−) containing 0.125% Trypsin (Thermo Fisher Scientific), 0.25 µL/mL DNase (Nippon Gene), 0.8 mM MgSO_4_ (KANTO), and 1.85 mM CaCl_2_ (KANTO) at 37°C for 15 min. The obtained cell suspension was passed through a 100 µm cell strainer (Falcon) and centrifuged with culture medium. After centrifugation, the cell pellet was resuspended in a culture medium and seeded on the cell culture plate. The culture medium was changed at 5 days in vitro (DIV). The cells were then used at DIV 8–9 for Aβ degradation assay and DIV 14–16 for qRT-PCR.

### Antibodies & Compounds

The following antibodies were used in this study: Anti-human Aβ, C99 (1:2500, 82E1, IBL, catalog #10323), Anti-GAPDH (1:1000, 6C5, Santa Cruz, catalog #sc-32233), Anti-Na^+^/K^+^-ATPase (1:1000, a6F, RRID: AB_528092, DSHB), Anti-APP (1:1000, APPc, IBL, catalog #8961), Anti-BACE1 (1:1000, BACE1c, Wako, catalog #18711), Anti-PS1 (1:1000, G1Nr5), Anti-sAPPβ (1:1000, 6A1, IBL, catalog #10321), Anti-IDE (1:1000, F-9, Santa Cruz, catalog #sc-393887), Anti-NEP (1:2000, R&D systems, catalog #AF1126). The following chemicals were used in this study: memantine (Daiichi-Sankyo Co. Ltd, stock solution was dissolved in distilled water, final concentration was 30 µM), L-glutamate (KANTO, catalog #17021-00, stock solution was dissolved in distilled water, final concentration was 100 µM), NMDA (Sigma-Aldrich, catalog #M3262, stock solution was dissolved in distilled water, final concentration was 100 µM), IKK-16 (Cayman chemical, catalog #13313, stock solution was dissolved in DMSO, final concentration was 2 µM for *in vitro* assay and 100 µM for *in vivo* assay), JSH-23 (Cayman chemical, catalog #15036, stock solution was dissolved in DMSO, final concentration was 30 µM), Trichostatin A (TSA) (Wako, catalog #203-17561, stock solution was dissolved in DMSO, final concentration was 1 µM), cOmplete^TM^ protease inhibitor cocktail (Roche Applied Science, catalog #11836145001), PhosSTOP^TM^ phosphatase inhibitor cocktail (Roche Applied Science, catalog #04906837001), Amyloid β-Protein (Human, 1-40) (Peptide Institute, catalog #4307-v, stock solution was dissolved in DMSO, final concentration was 20 nM).

### Aβ degradation assay

H4 cells and DIV 8–9 primary astrocytes were treated with synthetic human Aβ40 and several compounds. After incubation, the levels of remaining Aβ40 in the conditioned medium were analyzed by immunoblotting. The incubation times were 24 hours for H4 cells and 48 hours for primary astrocytes. For the input samples, fresh culture medium with synthetic human Aβ40 was prepared and incubated for the same durations.

### qRT-PCR

Total RNA was extracted from cells and hippocampal samples using ISOGEN (Nippon Gene, catalog #311-02501). cDNA was synthesized from purified RNA using the ReverTra Ace^®^ qPCR RT Master Mix with gDNA Remover (TOYOBO, catalog #FSQ-301). cDNA was mixed with gene-specific primers and THUNDERBIRD^®^ Next SYBR^TM^ qPCR Mix (TOYOBO, catalog #QPS-201) and then analyzed using a LightCycler^®^ 480 Instrument II (Roche). The PCR conditions were as follows: 1 cycle of pre-incubation (95°C, 1 min), followed by 50 cycles of denaturation (95°C, 10 sec), annealing (60°C, 30 sec), and extension (72°C, 1 min).

Specificity was confirmed by melting curve analysis and agarose gel DNA electrophoresis. The following primer pairs were used: 5’-GCACCGTCAAGGCTGAGAAC-3 (forward) and 5’-TGGTGAAGACGCCAGTGGA-3’ (reverse) for *GAPDH*; 5’-CGAGCCCAGATGTGACCTTT-3 (forward) and 5’-GTCACCATTGCAGGCGTTTT-3’ (reverse) for *KLK7*; 5’-AACGACCCCTTCATTGAC-3’ (forward) and 5’-GAAGACACCAGTAGACTCCAC-3’ (reverse) for *Gapdh*; 5’-AAGGACCTGCTGGGGAAAAC-3’ (forward) and 5’-CCCCTGAGTCACCATTGCAC-3’ (reverse) for *Klk7*. The amount of target mRNA was normalized to *GAPDH/Gapdh* mRNA in each sample, and the expression level of the indicated target gene was evaluated as the relative expression level.

### Luciferase reporter assay

The 238-bp promoter region of the human *KLK7* gene was amplified from genomic DNA by PCR using the following primers: 5’-TTTGCTAGCGACTGTGGGACCAGAATGTGCG-3’ (forward) and 5’-TTTAAGCTTTCTGATGTGATCCAAGTTCCGACTTG-3’ (reverse). The amplified human *KLK7* promoter and pGL4.16 [luc2CP/Hygro] firefly luciferase vector (Promega, catalog #E671A) were digested by Nhel and HindIII. After digestion, these constructs were gel-purified and ligated using Ligation high Ver.2 (TOYOBO, catalog #LGK-201). The luciferase constructs containing either a deletion mutant or a motif mutant of the κB motif located in the 238-bp promoter region were generated by PCR using the following primers: 5’-AACCGCAAATGCAGGGTCGGCTCCGCCTGCACCC-3’ (forward) and 5’-GCAGGCGGAGCCGACCCTGCATTTGCGGTTCTGG-3’ (reverse) for the deletion mutant; 5’-AACCGCAAATGCAGGCGCCCGGGACCCAGAGTCGGCTCCGCCTGCACCC-3’ (forward) and 5’-GCAGGCGGAGCCGACTCTGGGTCCCGGGCGCCTGCATTTGCGGTTCTGG-3’ (reverse) for the motif mutant. For transient expression of the luciferase constructs in H4 cells, Lipofectamine^®^ LTX Reagent & Plus™ Reagent (Thermo Fisher Scientific, catalog #15338100) was used following the manufacturer’s protocol. A Renilla luciferase vector, pRL-TK (Promega, catalog #E2241), was co-transfected as an internal control for transfection efficiency. For the generation of H4 stable cell lines expressing the luciferase constructs, polyethylenimine was used for transfection, followed by selection with Hygromycin B. For the luciferase reporter assay, cells were seeded and cultured for 3 days. Cells were then treated with compounds and collected 24 hours after treatment for luminescence detection. Luminescence was generated using the PicaGene^®^ Dual SeaPansy Luminescence Kit (TOYO B-NET, catalog #PD-11) and measured with the Mithras LB940 (BERTHOLD). In the analysis of transiently expressed luciferase constructs,

Firefly luciferase activity was normalized to Renilla luciferase activity. For the stable cell line analysis, Firefly luciferase activity was normalized to total protein concentration, determined using the Pierce^TM^ BCA Protein Assay Kit (Thermo Fisher Scientific).

### Alamar blue assay

Cell viability of H4 cells and primary astrocytes was measured using the alamarBlue™ Cell Viability Reagent (Thermo Fisher Scientific, catalog #DAL1025). After treatment with various compounds, cells were incubated for 2 hours in DMEM containing 10% Alamar Blue, and the fluorescence intensity of the conditioned medium was measured using a SpectraMax^®^ iD3 (MOLECULAR DEVICES).

### Animals

*App^NL-G-F/NL-G-F^* mice were originally generated at RIKEN Brain Science Institute, Japan^29^. *Klk7*^−/−^ mice was generated as previously described^17^. Briefly, sperm from *Klk7^tm1^*(KOMP)*^Vlcg/+^* heterozygous mice (project ID: VG14816) was generated through the trans-NIH Knockout Mouse Project [VelociGene at Regeneron Inc. (U01HG004085) and the CSD Consortium (U01HG004080)] and obtained from the KOMP Repository at UC Davis and CHORI (U42RR024244) (www.komp.org). *In vitro* fertilization using this sperm was performed at the Center for Disease Biology and Integrated Medicine, Graduate School of Medicine, The University of Tokyo. Genotypes were determined by PCR analysis of genomic DNA extracted from tail samples using alkali extraction. The following primers were used: 5’-CAGAGTGCCCAGAAGATCAAGG-3’ (Reg-Klk7-wtF) and 5’-CTGAGGCAATCTCACCGTCTGG-3’ (Reg-Klk7-wtR) for *Klk7* wild-type locus; 5’-GCAGCCTCTGTTCCACATACACTTCA-3’ (Reg-Neo-F) and 5’-ACCACACAACAACAGTCTCTCTTGC-3’ (Reg-Klk7-R) for *Klk7* knockout locus; 5’-ATCTCGGAAGTGAAGATG-3’ (E16WT) and 5’-TGTAGATGAGAACTTAAC-3’ (WT) for *App* wild-type locus; 5’-ATCTCGGAAGTGAATCTA-3’ (E16MT) and 5’-CGTATAATGTATGCTATACGAAG-3’ (loxP) for *App* knock-in locus.

### Injection of IKK-16 into the mouse hippocampus

Injection of IKK-16 was performed as previously described^60^. Briefly, 4 µL of 100 µM IKK-16 and 1% DMSO (Vehicle) in PBS were injected at 0.3 µL/min into the left and right sides of the hippocampus (anterior-posterior: −2.0 mm from bregma, medial-lateral: ±1.5 mm from midline, dorsal-ventral: 1.7 mm below brain surface) of 14–16-month-old female *App^NL-G-F/NL-G-F^*, *App^NL-G-F/NL-G-F^*; *Klk7*^−/−^ mice, and 7-month-old male wild-type mice. Hippocampal samples were collected from these mice 24 hours after injection for qRT-PCR analysis and 48 hours after injection for immunoblotting analysis.

### Brain extraction and fractionation

Hippocampal samples were homogenized in Tris buffer (50 mM Tris-HCl pH 7.6, 150 mM NaCl, cOmplete^TM^ protease inhibitor cocktail, and PhosSTOP^TM^ phosphatase inhibitor cocktail) using a mechanical homogenizer and centrifuged at 200,000×g for 20 min at 4°C. The supernatant was collected as the Tris buffer-soluble fraction (TS-soluble fraction), which mainly contains soluble proteins. The pellets were then homogenized in 2% Triton X-100/Tris buffer, centrifuged at 200,000×g for 20 min at 4°C, and the supernatant was collected as the Triton X-100-soluble fraction, which mainly contains membrane proteins. The pellets were then homogenized in 2% SDS/Tris buffer at room temperature, incubated at 37°C for 1 hour, and centrifuged at 200,000×g for 20 min at 20°C. The pellets were then sonicated with 70% formic acid and centrifuged at 200,000×g for 20 min at 4°C. The supernatant was freeze-dried for 2 hours using a Savant^TM^ SPD131DDA SpeedVac^TM^ Concentrator (Thermo Fisher Scientific), and the pellets were dissolved in DMSO as the formic acid fraction (SDS-insoluble fraction), which mainly contains insoluble proteins. Protein concentrations of the TS-soluble fraction and Triton X-100-soluble fraction were determined using the Pierce^TM^ BCA Protein Assay Kit (Thermo Fisher Scientific).

### Immunoblotting

Samples were dissolved in Laemmli sample buffer (2% SDS, 80 mM Tris-HCl pH 6.8, 10% glycerol, 0.0025% Brilliant green (Wako), and 0.00625% Coomassie Brilliant Blue G250 (Nacalai Tesque)) and boiled for 5 min at 100°C. For the detection of membrane proteins in 2% Triton X-100-soluble fraction extracted from mouse hippocampus, samples were incubated for 5 min at 65°C instead of boiling. Immunoblotting was performed as previously described^60^. Immunoreactivity was assessed by detecting the chemiluminescence of the samples. Chemiluminescence was generated using the ImmunoStar™ Reagents (Wako, catalog #291-55203) and detected with the ImageQuant™ LAS 4000 (GE Healthcare). The amount of target protein in the TS-soluble fraction was normalized to GAPDH, while the amount of target protein in the Triton X-100-soluble and SDS-insoluble fractions was normalized to Na^+^/K^+^-ATPase. The expression level of the indicated target protein was evaluated as the relative expression level.

### Data availability

Two public RNA-seq datasets were obtained from the AMP-AD Knowledge Portal (https://www.synapse.org/#!Synapse:syn2580853) as previously described^60,61^: the Mayo RNA-seq^27^ and MSBB studies^28^. The Mayo RNA-seq study includes temporal cortex samples from 164 subjects, comprising control (n=80) and AD (n=84) subjects. We assessed the expression levels of *KLK7*, *RELA*, *RELB*, *REL, NFKB1*, and *NFKB2* between control and AD subjects using a simple model (syn6090804), adjusting for key covariates: age at death, gender, RNA integrity number (RIN), source, and flow cell. For the MSBB study, we obtained clinical information of each subject, RNA-seq covariates, and normalized RNA read counts of above targets (syn7391833). As described in the previous report (syn20801188), gene level expression (read counts) was corrected for known covariates factors, including postmortem interval, race, batch, sex, RIN and exonic rate to remove the confounding effects. The trimmed mean of M values normalization method was used to estimate scaling factors and adjust for differences in library sizes. We selected 201 samples of the parahippocampal gyrus (Brodmann area 36) from subjects and excluded the samples without the information of the Braak NFT stage. These data were applied and analyzed using Python Jupyter Notebook. We compared the mRNA expression levels of the indicated target genes between healthy control subjects (CT) and AD patients. CT subjects were defined as those with an NP.1 stage, a neuropathology category measured by CERAD, equal to 1 (n=64), while AD patients were defined as those with stages ranging from 2 to 4 (n=137).

### Statistical analysis

Data are presented as mean values, and error bars indicate standard error of the mean (SEM). For data analysis, Student’s *t*-test, paired *t*-test, Dunnett’s test, and Bonferroni’s multiple comparison test were performed using GraphPad Prism 6. For the analysis of public RNA-seq datasets, Mann–Whitney *U* test was performed to compare gene expression levels, and Kendall rank correlation coefficient was performed to assess the correlation levels. A p-value<0.05 was considered to have a significant difference.

### Ethical approval

All experiments using animals in this study were performed according to the guidelines provided by the Institutional Animal Care Committee of the Graduate School of Pharmaceutical Sciences at the University of Tokyo (Protocol P4-29).

## Acknowledgements

This work was supported by Japan Society for the Promotion of Science (JSPS) Grants-in-aid for Scientific Research (A) 19H01015 and 23H00394 to T. Tomita; Grant-in-Aid for Transformative Research Areas (B) 22H05036 to Y. Hori; Japan Agency for Medical Research and Development Strategic Research Program for Brain Sciences 19dm0107056h0004 to T. Tomita; Japan Science and Technology Agency Moonshot R&D under grant number JPMJMS2024 to T. Tomita. C-J. Sung is a research fellow of the JSPS (24KJ0807). K. Kikuchi was a research fellow of the JSPS (18J14940). C-J. Sung and Y-W. Chiu were scholarship students of Japan-Taiwan Exchange Association.

The results of the MayoRNAseq Study and Mount Sinai Brain Bank data were published in whole or in part based on data obtained from the AMP-AD Knowledge Portal (https://adknowledgeportal.synapse.org/). The MayoRNAseq Study data were provided by the following sources: Mayo Clinic Alzheimer’s Disease Genetic Studies, led by Dr. Nilufer Taner and Dr. Steven G. Younkin (Mayo Clinic, Jacksonville, FL) using samples from the Mayo Clinic Study of Aging, the Mayo Clinic Alzheimer’s Disease Research Center, and the Mayo Clinic Brain Bank. Data collection was supported through funding by National Institute on Aging Grants P50 AG016574, R01 AG032990, U01 AG046139, R01 AG018023, U01 AG006576, U01 AG006786, R01 AG025711, R01 AG017216, and R01 AG003949; National Institute of Neurological Disorders and Stroke Grant R01 NS080820; CurePSP Foundation; and Mayo Foundation. Study data include samples collected through the Sun Health Research Institute Brain and Body Donation Program (Sun City, AZ). The Brain and Body Donation Program was supported by National Institute of Neurological Disorders and Stroke (U24 NS072026 National Brain and Tissue Resource for Parkinson’s Disease and Related Disorders), National Institute on Aging (P30 AG19610 Arizona Alzheimer’s Disease Core Center), Arizona Department of Health Services (Contract 211002, Arizona Alzheimer’s Research Center), Arizona Biomedical Research Commission (Contracts 4001, 0011, 05-901, and 1001 to the Arizona Parkinson’s Disease Consortium), and Michael J. Fox Foundation for Parkinson’s Research. The Mount Sinai Brain Bank data were generated from postmortem brain tissue collected through the Mount Sinai Veterans Administration Medical Center Brain Bank and were provided by Dr. Eric Schadt (Mount Sinai School of Medicine).

We thank Dr. Takaomi C. Saido (RIKEN Center for Brain Science) and Dr. Takashi Saito (Nagoya City University) for providing *App^NL-G-F/NL-G-F^* mice, as well as our current and former laboratory members for valuable discussions.

## Author contributions

Y. Sudo, K. Kikuchi, and T. Tomita designed research; Y. Sudo and C-J. Sung performed experiments; Y. Sudo, C-J. Sung, Y-W. Chiu, and T. Tomita analyzed data; Y. Sudo, C-J. Sung, and T. Tomita wrote the paper; Y. Sudo, C-J. Sung, S. Takatori, Y. Hori, and T. Tomita edited the paper. All authors discussed the results and approved the final manuscript.

